# Understanding how an amphicarpic species with a mixed mating system responds to fire: a population genetic approach

**DOI:** 10.1101/2020.10.18.344036

**Authors:** Elena M. Meyer, Joel F. Swift, Burgund Bassüner, Stacy A. Smith, Eric S. Menges, Brad Oberle, Christine E. Edwards

## Abstract

Amphicarphic species produce both aboveground and belowground seeds; the belowground seeds have been proposed to be an adaptation to disturbed sites because they are protected belowground, enabling them to persist and recolonize a site after disturbance. However, it is unknown whether such seeds indeed serve as the main colonizers after a disturbance. The amphicarpic species *Polygala lewtonii* is endemic to fire-prone Florida sandhill and scrub and is among only a few species with three flower types (aboveground chasmogamous flowers and both above and belowground cleistogamous flowers). The goal of this study was to understand whether recolonization of sites by *P. lewtonii* was accomplished primarily through germination of belowground seed. First, we quantified the outcrossing rate in seeds produced by aboveground chasmogamous flowers to determine whether we could detect differences in colonization of between seeds produced aboveground vs. belowground. Approximately 25% of seeds from aboveground chasmogamous flowers showed evidence of cross pollination and the seeds showed greater heterozygosity and lower inbreeding coefficients than pure selfing, indicating that it is possible to differentiate between selfed and non-selfed seed types in postfire colonization. Second, we analyzed genetic diversity, inbreeding, and genetic structure of the populations before and after a prescribed fire. If heterozygosity and admixture increased, and spatial population genetic structure and inbreeding decreased, this would indicate that fire promoted germination of outcrossed seed from aboveground flowers. However, inbreeding increased and spatial genetic structure and admixture decreased after fire, suggesting that selfed seed produced by belowground flowers predominantly recolonized the site after fire. Thus, amphicarpy is a powerful adaptation to fire-maintained environments by producing seeds that are well suited to the range of conditions presented by a highly variable, disturbance prone habitat.

## Introduction

Understanding how reproductive strategies evolve in disturbance-maintained ecosystems connects basic evolutionary theory to conservation applications. Episodic mortality from regular disturbance tends to reduce population genetic variation and increase inbreeding (Dolan et al, 2009; Davies et al, 2016), while selecting for life history traits that maintain reproductive potential (Lytle 2001). Associated reproductive strategies, like self-compatibility or seed dormancy, may promote reproductive assurance while constraining genetic variation and dispersal to the point of increasing extinction risk (Goldberg and Igić 2012, Vamosi et al, 2018). Consequently, effective conservation management of species endemic to disturbance-maintained ecosystems requires special attention to connections between disturbance, demography, mating system and genetic diversity.

A relatively common reproductive strategy among plant species in disturbance-prone habitats is a mixed mating system. Mixed mating is characterized by 1) both self- and cross-fertilization, 2) a population outcrossing rate that departs significantly from both zero and one, and 3) a proportion of selfing between 0.2 < *t* ≤ 0.8 (Goodwille et al, 2005). Mixed mating systems have both drawbacks and advantages. They are hypothesized to be an adaptation to ecosystems that regularly experience disturbance by providing reproductive assurance through self-fertilization when pollinators or conspecifics are absent (Cheplick 1987; Symonides, 1988). The most obvious drawback is that reliance on self-fertilization may lead to inbreeding depression (i.e., decreased fitness of individuals because of increased homozygosity for recessive deleterious alleles); however, the purging of deleterious alleles through selection may mitigate this effect by becoming more effective in heavily selfing lineages (i.e., “selfers”) (Holsinger, 1988).

A second, less common life history trait proposed to be an adaptation to frequent disturbance is amphicarpy, a form of dimorphism in which both aerial and subterranean flowers and seeds are produced on the same individual (Cheplick, 1987). Amphicarpic plants often exhibit seed polymorphism (Symonides, 1988), with belowground, obligately selfing, or cleistogamous (CL) flowers producing seeds that are typically larger but with a limited dispersal potential, and aboveground, open-pollinated, or chasmogamous (CH) flowers producing smaller seeds that are suitable for dispersal across larger distances (Cheplick, 1987; Kaul, 2000; Zhang et al 2020). The belowground seeds were proposed to be an adaptation to disturbance-prone habitats, as belowground seeds remain below the soil surface, are protected from disturbances such as fire, and may be able to quickly recolonize a site after a disturbance provided that they remain viable and maintain dormancy until stimulated by disturbance (Cheplick and Quinn 1987, Cheplick and Quinn 1988). Although previous studies found that seeds buried belowground often showed greater germination after a fire (Cheplick and Quinn 1987, Cheplick and Quinn 1988), few studies have empirically tested some of the basic assumptions of the hypothesis that amphicarpy is an adaptation to disturbance, such as whether recolonization by amphicarpic species after a disturbance is primarily accomplished through germination of seeds produced by belowground flowers.

An ideal species for investigating whether amphicarpy is an adaptation to disturbance is *Polygala lewtonii* Small (Polygalaceae), or Lewton’s polygala, a federally endangered (USFWS, 1999), amphicarpic plant species. *P. lewtonii* is endemic to Florida sandhill and scrub, which are fire-maintained ecosystems. Previous studies found evidence that *P. lewtonii* is adapted to fire; smoke stimulated seed germination (Lindon and Menges, 2008) and population sizes increased dramatically after burning (Weekley and Menges, 2012). This species is one of only a few that exhibits mixed mating and amphicarpy via three types of flowers: 1) above-ground CH flowers, 2) aboveground CL flowers, and 3) belowground CL flowers. In *P. lewtonii*, open-pollinated CH flowers also have a delayed selfing mechanism (Weekly and Brothers, 2005), which was proposed to provide reproductive assurance in the absence of pollinators (Cheplick 1987; Symonides, 1988). However, the reproductive assurance provided by this mechanism was shown to be limited for *P. lewtonii*, as fruit production dropped markedly when pollinators were excluded (Weekly and Brothers, 2005). Although *P. lewtonii* matures both aboveground and belowground fruits, the different types of flowers vary in rates of fruit production, with CH flowers producing over seven times the number of fruits produced by above- and below-ground cleistogamous flowers combined (Koontz et al., 2017). All seeds produce an elaiosome, a fleshy appendage that attracts ants for seed dispersal (Zomlefer, 1989). Ants have been observed to rigorously collect the aboveground seeds (USFWS, 2009; Menges and Weekly, 2003) and may disperse them up to several meters. However, there is no evidence that ants access or disperse the belowground CL seeds (Weekly and Brothers, 2006), meaning that their dispersal is likely limited by the length of the rhizomes (<1m).

Although *P. lewtonii* was the subject of a previous fine-scale population genetic analysis to understand its predominant mating system and patterns of population genetic structure, it is unknown how these factors are affected by fire. The previous population genetic study in *P. lewtonii* found most reproduction occurred via inbreeding, that outcrossing rates were low, and that genetic variation was structured at a very fine scale (Swift et al 2016). The low outcrossing rate was unexpected because the showy, open-pollinated CH flowers have high rates of fruit set, even when pollinators were excluded (71.2 %; Weekly and Brothers, 2006). Swift et al. (2016) hypothesized that the low outcrossing rate may be due to a lack of recent fire, which may be necessary to stimulate germination of the outcrossed seed produced aboveground. If this is the case, then we would expect to see an increase in heterozygosity and a decrease in inbreeding and genetic structure after a fire. Conversely, other authors have proposed that belowground CL flowers and seeds of amphicarpic species may serve to re-colonize a site after a disturbance (Cheplick et al, 1987, Cheplick et al 1988), in which case we would expect an increase in inbreeding and genetic structure after a fire. A study to investigate how the outcrossing rate and patterns of genetic structure change in response to fire is needed to address this question, with important implications for understanding the ecological and evolutionary function of amphicarpy.

In this study, we used a population genetic approach to analyze the dynamics of recolonization of the amphicarpic species *P. lewtonii* after fire. Our analyses employed two experiments. The first tested whether we could differentiate among seed types in postfire recolonization. We quantified the outcrossing rate in seeds produced by aboveground CH flowers by comparing their genotypes to those of their maternal parents; we then tested whether they deviated from the patterns expected for 100% selfing of CL flowers. Because CH flowers represent a substantial investment and exhibited low fruit set when pollinators were limited (Weekly and Brothers, 2005), we hypothesized that a majority of seeds produced by aboveground CH flowers would be the product of outcrossing. Specifically, we expected to encounter non-maternal alleles that could not be the product of selfing in a majority of seeds and overall heterozygosity rates significantly exceeding the 50% reduction expected to accompany a single generation of pure selfing.

Second, we used a population genetic approach to compare the genetic diversity and structure of populations before and after fire. We tested how inbreeding coefficients, genetic diversity, and genetic structure changed after a prescribed burn in a natural population of *P. lewtonii*. If we found increased heterozygosity and admixture and decreased inbreeding and spatial population genetic structure after a prescribed fire, then this would support the hypothesis that burning stimulated the germination of seed from aboveground flowers. This assumes that a moderate proportion of aboveground seed is produced through outcrossing in CH flowers and that the relative contribution of aboveground CL flowers to the total aboveground seed production is small, as was shown previously (Koontz et al. 2017). Conversely, if we found decreased heterozygosity and admixture and increased inbreeding and among-group spatial population genetic structure after a prescribed fire, then this would support the hypothesis that burning stimulates germination of seed produced by belowground CL flowers, as they are obligately selfing and disperse very short distances. The results of these experiments have important consequences for the conservation of *P. lewtonii* and more broadly for our understanding of both the ecological and evolutionary function of amphicarpy.

## Materials and Methods

### Study Species

*Polygala lewtonii* (Polygalaceae) is a short-lived (2-10 years) perennial, reaching around 20 cm tall (Weekly and Brothers, 2006). The plant bears clusters of spatulate to linear-spatulate leaves that overlap on the glabrous stem like shingles (Small, 1933, USFWS, 1999). The aboveground CH flowers of the plant are deep pink to purple and occur on densely flowered terminal racemes (Weekly and Brothers 2006). A variety of insect species have been observed visiting the species, including sulfur butterflies and bee flies (Weekly and Brothers, 2006). In addition to the aboveground CH flowers, belowground white CL flowers occur sparsely on the rhizomes, and, rarely, solitary pale pink aboveground CL flowers occur in the axils of the aboveground leaves on short leafless branches (Wunderlin and Hansen, 2016; Weekly and Brothers, 2006). *P. lewtonii* flowers throughout most of the year, with temporal variation in the presence of each flower type; CH flowers are present in January–May, aerial CL flowers are present from June–January, and belowground CL flowers appear from July–February (Koontz et al, 2015). The fruit is oblong, ca. 4 mm long (Wunderlin and Hansen, 2016), dehiscent, and two-seeded (Zomlefer, 1989). The ellipsoid-cylindric seeds (Wunderlin and Hansen, 2016) bear stiff hairs and two aril-like outgrowths at the micropyle, referred to as elaiosomes, which attract ants that disperse only the aboveground seed (Zomlefer, 1989, Weekly and Brothers, 2006).

*Polygala lewtonii* is endemic to six counties in central Florida (Brevard, Highlands, Lake, Marion, Polk, and Osceola counties; Wunderlin et al, 2017), and occurs only in sandhill and scrub habitats on the Lake Wales and the Mount Dora Ridges in peninsular Florida (Menges et al, 2007). These xeric ecosystems experience fire as their dominant mode of ecological disturbance (Menges 2007), with more intense fires in scrub and more frequent fires in sandhill. In scrub, fire significantly affects the habitat structure by increasing bare ground and decreasing canopy cover, thus maintaining the ecosystem as a scrubland (Ashton, K. G., & Knipps, 2011).

Fire also changes the species composition of Florida scrub, increasing the frequency and abundance of scrub herbs because of post-burn reductions in shrubs, litter, and lichens (Weekly and Menges, 2003). Fire has a positive effect on populations of rare, endemic scrub plants, including increasing flowering and seedling recruitment (Slapcinsky et al, 2010). In sandhill, fire promotes species diversity and temporarily reduces cover (Heuberger & Putz 2003; Reinhart & Menges, 2004). Frequent fires in sandhill can increase the cover of grasses and reduce shrub cover (Reinhart and Menges 2004). The xeric uplands in central Florida have suffered from habitat loss and fragmentation (Weekley et al. 2008), as well as anthropogenic alteration of disturbance regimes, which include increased fire-return intervals and limited use of prescribed fires in land management (USFWS, 1999). Fire suppression is one of the most common and most serious threats to both Florida sandhill and scrub (Menges 2007).

### Sample collection

To understand the relative outcrossing rates in aboveground CH flowers, five seeds per plant were collected from the aboveground, open-pollinated flowers of ten maternal plants at the Carter Creek Tract of the Lake Wales Ridge Wildlife and Environmental Area (LWRWEA). We also sampled leaf tissue from the same plants for comparison with seed samples. Seeds were stored in paper envelopes at room temperature and leaf tissue was stored in silica gel desiccant until DNA extraction.

To understand the dynamics of recolonization after fire, individuals were initially sampled from the Carter Creek Tract of the LWRWEA in 2014, as described in Swift et al. (2016). The study area was initially defined in 2014 by compiling all known locations (GPS points) for *P. lewtonii* within the site, yielding an estimated range of ∼0.28 km^2^. Within this study area, four blocks were created, and within each block, eight collection plots were established that ranged from 1–4 m in radius containing a minimum of nine *P. lewtonii* individuals, with a minimum distance of 10 m separating collection plots in the same block. The closest plots in neighboring blocks were separated by larger distances (mean = 350 m, range 140–630 m between neighboring blocks). By design, these plots were placed at a very fine spatial scale to understand how genotypes and genetic clusters were distributed spatially, with the goal of providing insight into the patterns of reproduction and the dispersal of selfed versus outcrossed seeds in the species. In the 2014 sampling, leaves were collected from nine individuals per plot in each of eight plots per block, totaling 72 individuals per block, or 288 individuals in total across the four blocks. The site was then subjected to a prescribed fire in 2016, which burned all blocks and plots except block 1 (plots 1-8 in Swift et al; 2016) and plots 17, 19, 20, and 24. In spring of 2017, we resampled the 20 fully burned plots, collecting leaf tissue from individuals germinating after the fire (Table 1). After fire, we increased sampling to 12 individuals per plot, totaling 240 individuals (Table 1, Figure 1). To ensure that the pre- and post-fire data sets were comparable, only the pre-fire data from the 20 plots that were completely burned were included in the present study, such that the pre-fire data set contained 180 individuals.

**Table 1.**
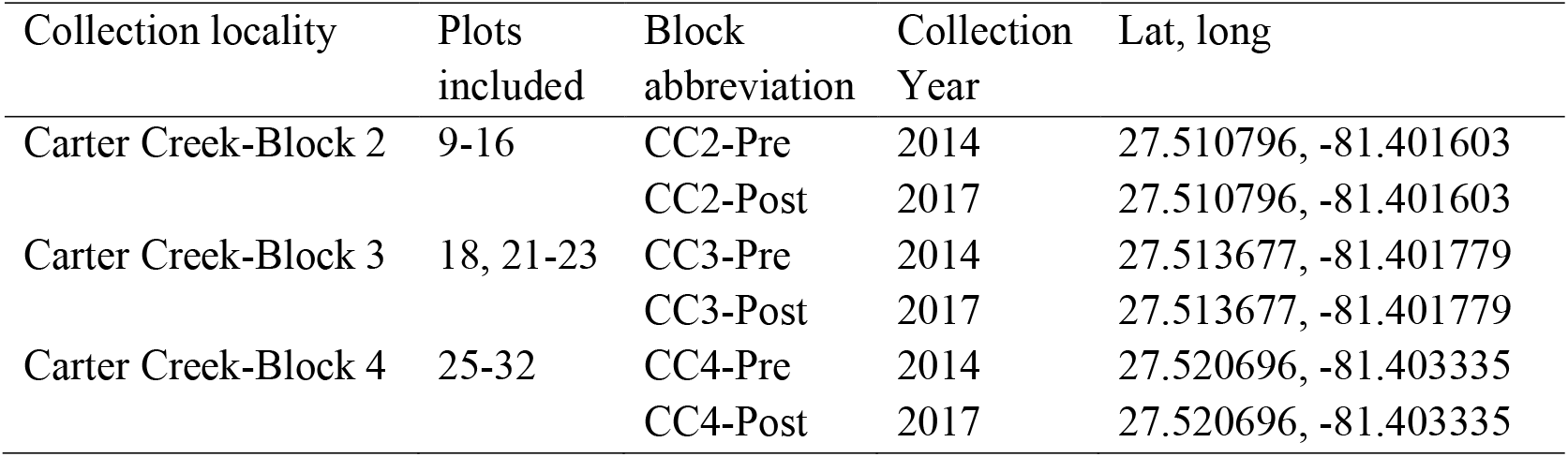
The block name, collection location, year, and latitude/longitude for pre- and post-fire sampling of *Polygala lewtonii*.

| Collection locality | Plots included | Block abbreviation | Collection Year | Lat, long |
| --- | --- | --- | --- | --- |
| Carter Creek-Block 2 | 9-16 | CC2-Pre | 2014 | 27.510796, -81.401603 |
|  |  | CC2-Post | 2017 | 27.510796, -81.401603 |
| Carter Creek-Block 3 | 18, 21-23 | CC3-Pre | 2014 | 27.513677, -81.401779 |
|  |  | CC3-Post | 2017 | 27.513677, -81.401779 |
| Carter Creek-Block 4 | 25-32 | CC4-Pre | 2014 | 27.520696, -81.403335 |
|  |  | CC4-Post | 2017 | 27.520696, -81.403335 |

**Figure 1.**
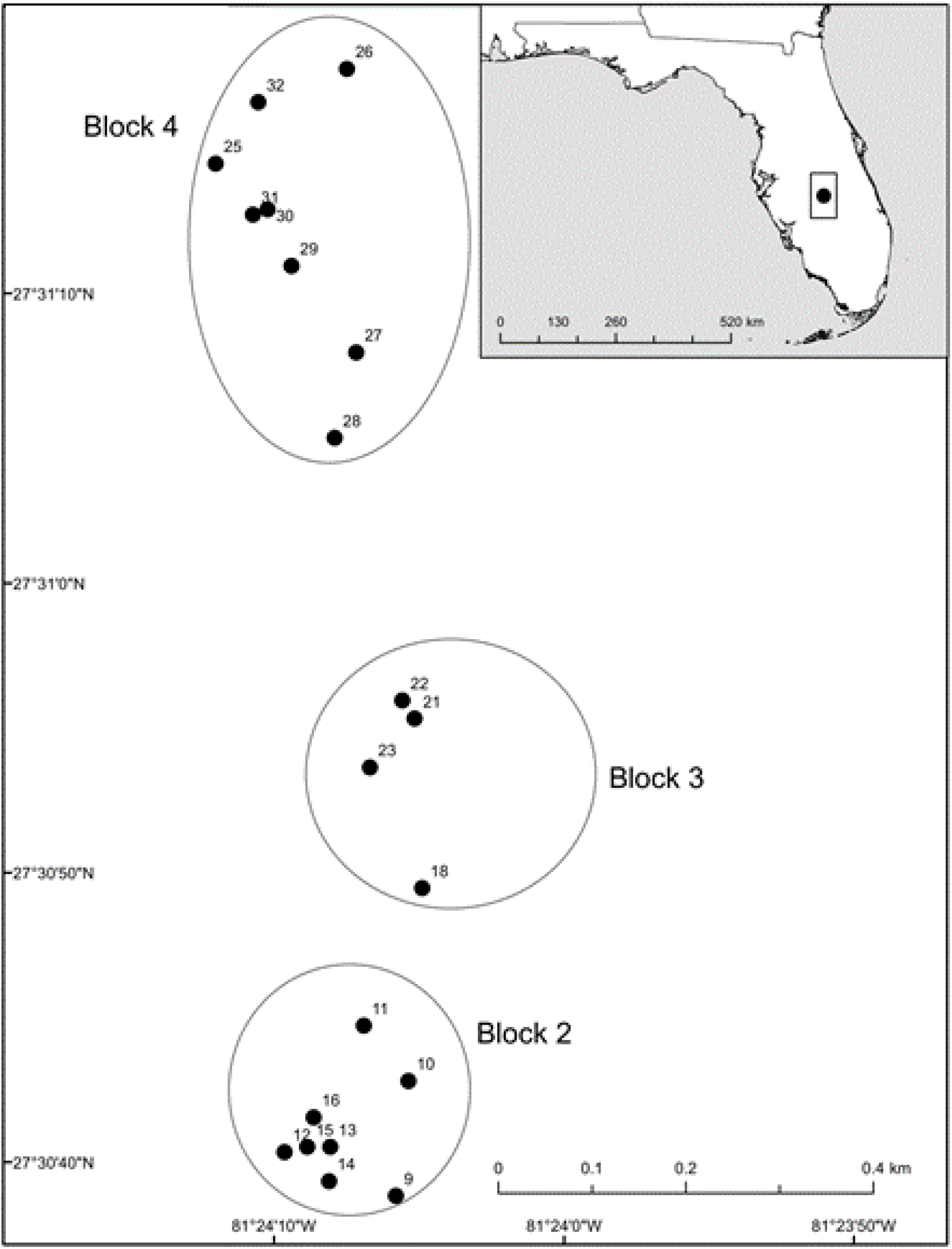
Plots (points) and blocks (circled) at Carter Creek Lake Wales Ridge National Wildlife Refuge in Central Florida. Only plots that were burned and re-sampled are shown.

### Microsatellite Analysis

DNA extractions were performed using a CTAB DNA Extraction Protocol (Doyle and Doyle 1987) that was modified by using smaller volumes, a smaller initial sample of plant tissue, and an additional wash step with cold 95% ethanol. The seed and leaf tissue samples were genotyped using 11 microsatellite loci as described in Swift et. al. (2016). Fragment analysis and scoring were carried out using previously developed automated fragment scoring panels in GeneMarker version 1.6 (Soft Genetics LLC), the results of which were then checked manually. Individuals with >50% missing data were removed from the analysis, resulting the removal of 10% of the individuals in the seed data set and ∼2% of individuals in the pre- and post-fire analysis.

### Data analysis of seed samples and simulation of mating system

The seed genotype data was initially compared to that of the maternal tissue. Genotyping error was identified if the seed contained no alleles from the maternal plant at a locus, in which case it was removed from the analysis. To analyze the extent to which seeds produced by aboveground CH flowers were derived from inbreeding vs. outcrossing, we counted the number of non-maternal alleles found in the offspring, as these alleles would presumably be contributed by the pollen donor. Seeds were placed into three categories: individuals having zero non-maternal alleles, those having only one non-material allele, and those having more than one non-maternal allele. Individuals with zero non-maternal alleles were products of either selfing or bi-parental inbreeding involving highly genetically similar individuals. Individuals with one non-maternal allele were either the product of cross-pollination between closely related parents or genotyping error. Individuals with multiple non-maternal alleles were likely products of outcrossing.

For seed data, to directly compare the heterozygosity of seeds from aboveground CH flowers to the expected results under pure selfing, we conducted a simulation of pure inbreeding in R version 3.4.1 (R Core Team 2013). Specifically, for each parent plant, we randomly sampled 10 alleles from each locus that we assembled into five multilocus selfed-seed genotypes. We then repeated the simulation of pure-selfing 1000 times. For each simulated seed dataset, we calculated five different indices of individual heterozygosity using the function GENHET (Coulon, 2009), including PHt, the proportion of heterozygous loci; Hs-observed and Hs-expected, which are standardized measures of observed and expected heterozygosity; IR, or internal relatedness, which weighs shared rare alleles more highly than shared common alleles; and HL, homozygosity by locus, which is expected to follow the opposite pattern of the other four metrics. The script used for this analysis is presented in Supporting Information–Appendix S1. We identified significant evidence for outcrossing when an observed heterozygosity metric exceeded the 95% quantile of metrics calculated for datasets simulating pure inbreeding (Figure S1).

### Analysis of changes in genetic diversity and inbreeding after fire

For each block both pre- and post-fire, we first used GenAlEx version 6.503 (Peakall et al, 2012) to calculate the average number of alleles per locus (*A*) and number of private alleles per population (A_P_). We used FSTAT v. 2.94 (Goudet 1995) to calculate allelic richness using rarefaction. We also placed individuals into groups containing all pre-fire or post-fire individuals and used GenAlEx to calculate the number of private alleles; emergence of new private alleles after fire reflects seedbank contributions by individuals not sampled during the original survey. We also used GenAlEx to calculate observed and expected heterozygosity (*H*_*O*_ and *H*_*E*_); if outcrossed seeds preferentially germinated in response to fire, then we would expect *H*_*O*_ to increase post-fire, whereas if seeds that germinated were predominantly produced by inbreeding, then we would expect *H*_*O*_ to decrease or possibly remain at pre-fire levels, as the species previously showed very low *H*_*O*_ values. To detect whether fire stimulated germination of selfed vs. outcrossed seeds, we also used GenAlEx to measure the inbreeding coefficient (*F*). Because technical artifacts such as genotyping errors and null alleles (i.e., undetected alleles that result from differential amplification success during PCR, (Chapuis and Estoup 2007) inflate inbreeding coefficients, we used the program INEST version 2 (Chybicki and Burczyk 2009) to calculated an alternative inbreeding coefficient which took null alleles and genotyping errors into account (*F*^*B*^). INEST uses a population inbreeding model to simultaneously measure the frequency of null alleles at each locus and calculate the inbreeding coefficient in each population. We used the Bayesian MCMC approach with 5,000,000 iterations, a burn-in of 500,000, and sampled every 250^th^ iteration to calculate a revised *F*^*B*^, which incorporated null alleles in each population. If fire stimulated the germination of seeds produced by inbreeding, then we would expect an increase in *F* and *F*^*B*^ post-fire, whereas if seed produced by outcrossing predominantly germinated post-fire, then we would expect a decrease in *F* and *F*^*B*^.

### Analysis of changes in genetic structure before and after fire

We also analyzed how patterns of genetic structure changed in response to fire. To understand how the hierarchical partitioning of genetic variation within and among populations changed over time, we conducted AMOVA analyses as implemented in GenAlEx. Analyses were conducted separately for pre-fire and post-fire samples, and we conducted two tests for each, with samples divided into either blocks or plots. If selfed belowground seeds were preferentially germinating in response to fire, then we would expect increased among-group variance after a fire and decreased within-group variance because the belowground seeds are obligately selfing and can disperse only across the length of a rhizome (∼1m). If outcrossed seed preferentially germinated, then we would expect among-group variation to decrease and within-group variation to increase, as cross pollination between plants in different blocks would contribute to a higher proportion of admixture in the offspring.

To estimate genetic structure independent of a-priori population designations, we utilized InStruct (Gao et al, 2007), which was specifically designed to measure patterns of genetic structure in species with inbreeding or selfing mating systems. It differs from STRUCTURE (Prichard et al, 2000) in that it eliminates the assumption of Hardy-Weinberg equilibrium within genetic clusters and jointly estimates the selfing rate and population structure. We analyzed patterns of genetic structure for a data set composed of both the pre- and post-fire data using the default settings in InStruct, except we varied the number of groups, *K*, from 1 to 15, employed 10 independent chains of the MCMC algorithm for each K, and used a burn-in of 500,000 iterations and a run length of 1,000,000 iterations for each chain. To ensure convergence and repeatability, we examined the groupings across all runs at each *K* in CLUMPAK (Kopelman et al. 2015, Gilbert et al. 2012). To determine the optimal value of *K*, we plotted the Deviance Information Criterion (Spiegelhalter et al, 2002) and the -ln likelihood values from InStruct and selected the value where they plateaued and showed clear patterns of genetic structure.

To quantify how admixture and genetic structure changed after a fire, for each plot both pre and post-fire, we inspected InStruct admixture proportions to identify the predominant genetic cluster to which individuals were assigned and calculated the percentage of individuals that showed their greatest assignment to the predominant cluster. We also calculated average admixture proportions at the predominant InStruct cluster across all individuals in a plot, both pre- and post-fire. If belowground, selfed seed predominantly germinated after a fire, then we would expect post-fire plots to show more individuals showing assignment to a predominant genetic cluster and greater average admixture proportions assigned to the predominant genetic cluster. If outcrossed seeds preferentially germinated after a fire, then we would expect post-fire plots to show fewer individuals showing assignment to a predominant genetic cluster and lower average admixture proportions assigned to the predominant genetic cluster.

Finally, to understand how grouping of individuals affects estimates of genetic diversity, structure, and admixture, we repeated analyses with individuals grouped into the predominant genetic cluster to which they were assigned by INSTRUCT. Analyses included estimates of genetic diversity, AMOVA, and average admixture proportions at the majority cluster.

## Results

### Outcrossing rate in aboveground CH flowers

Seeds produced by aboveground CH Flowers showed evidence of outcrossing. After removing 10 seeds from the analysis because of genotyping failure or genotyping errors, the final data set contained 40 seeds from nine maternal individuals, of which 30 out of 40 (75%) had only maternal alleles present, 4 of 40 (10%) differed from their mother by only one allele, and 6 of 40 (15%) differed from their mother by three or more alleles. When comparing observed inbreeding metrics to those simulated under 100% selfing, the internal relatedness among seeds from open-pollinated flowers (0.718) was well below the range of values simulated for pure selfing (0.900-0.985, Figure 2). Other heterozygosity metrics (PHt, HL, Hs-observed and expected) showed the same pattern (Supporting Information–Table S1).

**Figure 2.**
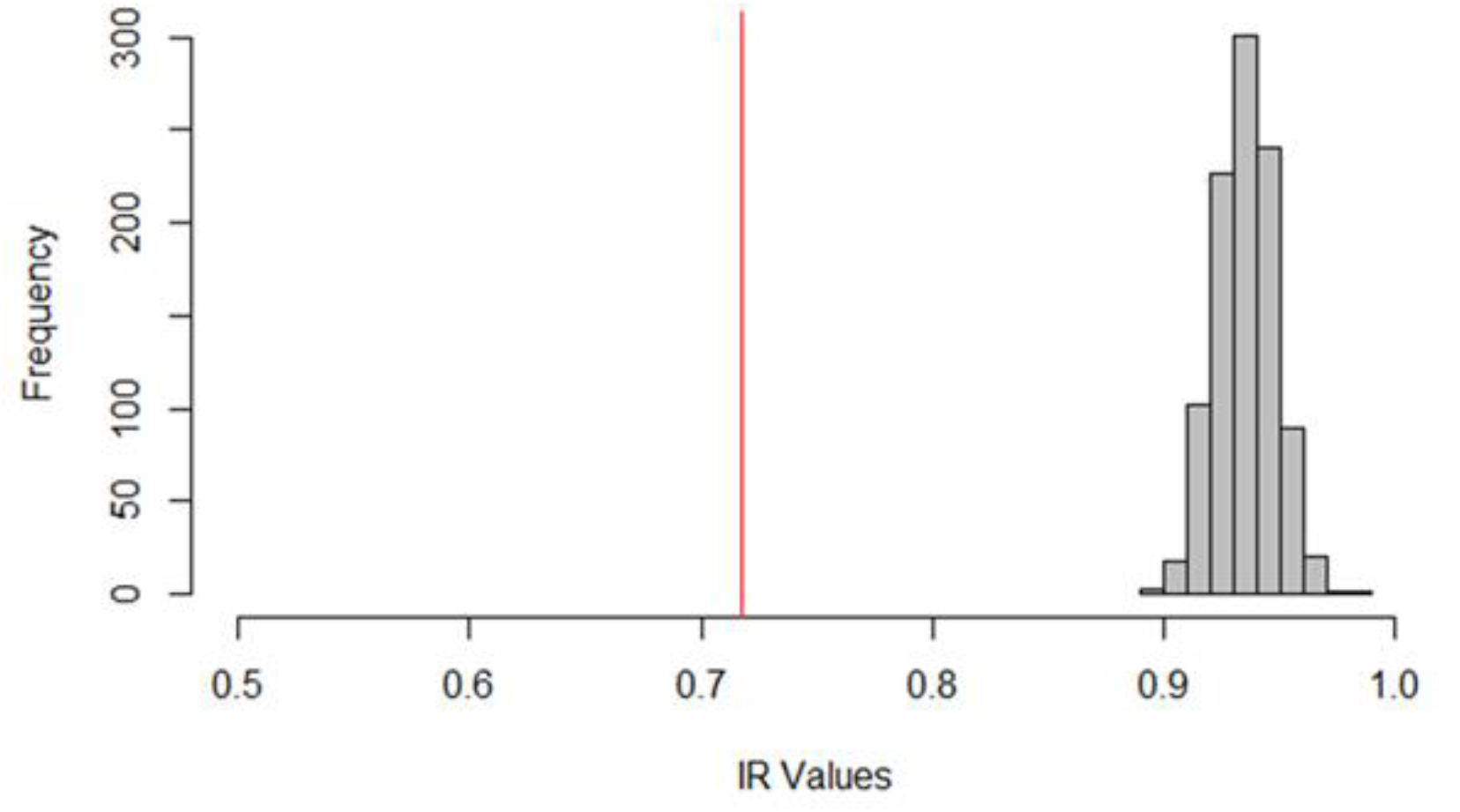
Observed values of heterozygosity in the seeds of chasmogamous flowers differ significantly from complete selfing simulations. Distribution of heterozygosity values measured by internal relatedness (IR) under simulated complete selfing (gray distribution) vs. the observed range of heterozygosity in seed samples (red line).

### Genetic diversity before and after fire

Burning had modest effects on genetic diversity. The mean number of alleles per population increased slightly after fire (mean A_Pre_ = 3.00, mean A_Post_ = 3.303). Analyses of allelic richness, which are corrected for differences in sample size, also showed a slight increase after fire (mean A_R-Pre_ = 2.71, mean A_R-Post =_ 2.79 Table 2). When grouped into blocks, the mean number of private alleles per block was 3.0 pre-fire and was 1.33 post-fire. When comparing private alleles between pre- and post-fire groups, seven alleles were unique to pre-fire populations, and eight alleles were unique post-fire (Supporting Information–Table S1). All eight post-fire private alleles were not observed previously, even when considering the whole range (Swift et al. 2016). All private alleles were present at low frequencies (range = 0.004-0.031%, mean = 0.01%; Supporting Information–Table S2), but together, alleles unique to one sampling time accounted for ∼17% of all alleles (Supporting Information–Table S2).

**Table 2.** The effects of fire on genetic diversity in *Polygala lewtonii*. Parameters include *N*, number of samples, *H*_*O*_, observed heterozygosity, *H*_*E*_, expected heterozygosity, *A*, average number of alleles per locus, *A*_*R*_ allelic richness based on a minimum sample size of 21 individuals, *Ap*, number of private alleles, *F*_*IS*_, inbreeding coefficient, and *F*_*IS*_^*B*^ the inbreeding coefficient taking null alleles into account,

| | $N$ | $H_O$ | $H_E$ | $A$ | $A_R$ | $A_P$ | $F_{IS}$ | $F_{IS}^B$ |
| --- | --- | --- | --- | --- | --- | --- | --- | --- |
| By block |  |  |  |  |  |  |  |  |
| CC2-Pre | 68 | 0.063 | 0.327 | 3.091 | 2.716 | 2 | 0.796 | 0.737 |
| CC3-Pre | 33 | 0.024 | 0.354 | 2.545 | 2.611 | 1 | 0.932 | 0.902 |
| CC4-Pre | 70 | 0.030 | 0.251 | 3.364 | 2.798 | 4 | 0.658 | 0.806 |
| Mean-Pre |  | 0.039 | 0.311 | 3.00 | 2.71 | 3.00 | 0.795 | 0.815 |
| CC2-Post | 96 | 0.077 | 0.34 | 3.727 | 2.612 | 3 | 0.733 | 0.765 |
| CC3-Post | 48 | 0.027 | 0.371 | 2.909 | 2.993 | 1 | 0.945 | 0.925 |
| CC4-Post | 96 | 0.038 | 0.255 | 3.273 | 2.773 | 0 | 0.757 | 0.846 |
| Mean-Post |  | 0.047 | 0.322 | 3.303 | 2.792 | 1.333 | 0.811 | 0.845 |

Values of expected heterozygosity (*H*_*E*_), observed heterozygosity (*H*_*O*_), and inbreeding coefficients increased slightly after fire. The mean *H*_*E*_ averaged across loci for pre-fire populations was 0.311, whereas it was 0.322 post-fire (Table 2). Values of also *H*_*O*_ increased slightly after burning (pre-fire mean=0.039, post-fire mean = 0.047) (Table 2). As found previously, mean values of *H*_*O*_ were much lower than expected heterozygosity both before and after fire, such that inbreeding coefficients (*F*) reached values close to one. Inbreeding coefficients ranged from 0.658 to 0.932 (mean = 0.795) in pre-fire populations and increased slightly post-fire, ranging from 0.733 to 0.945 (mean = 0.811) (Table 2). After correction for null alleles using INEST (Chybicki and Burczyk, 2009), *F*^*B*^ ranged from 0.737 to 0.902 (mean = 0.815) in pre-fire blocks and was greater in post-fire blocks, ranging from 0.765 to 0.925 (mean = 0.845) (Table 2).

### Patterns of genetic structure among blocks before and after fire

Burning also changed the hierarchical partitioning of genetic variation. Pre-fire samples had 26% of the variation partitioned among blocks, 68% among individuals within blocks, and 7% within individuals (Table 3). Post-fire, a greater percentage of variation was found among blocks (33%) and less was found among individuals within blocks (57%). Similar changes occurred at the relatively fine-scale sampling unit at the plot level (where plots are 1–4 m in radius, as compared to the larger blocks). Pre-fire samples at the plot level showed 46% of the variation partitioned among plots, 47% among individuals within plots, and 7% within individuals, whereas post-fire samples at the plot level showed 58% of the variation partitioned among plots, 31% among individuals within plots, and 11% within individuals (Table 3). F-statistics (F_ST,_ F_IS_ and F_IT_) derived from AMOVA also calculated These show a pattern of and an increased F_ST_ and decreased F_IS_ after fire (Table 3).

**Table 3.**
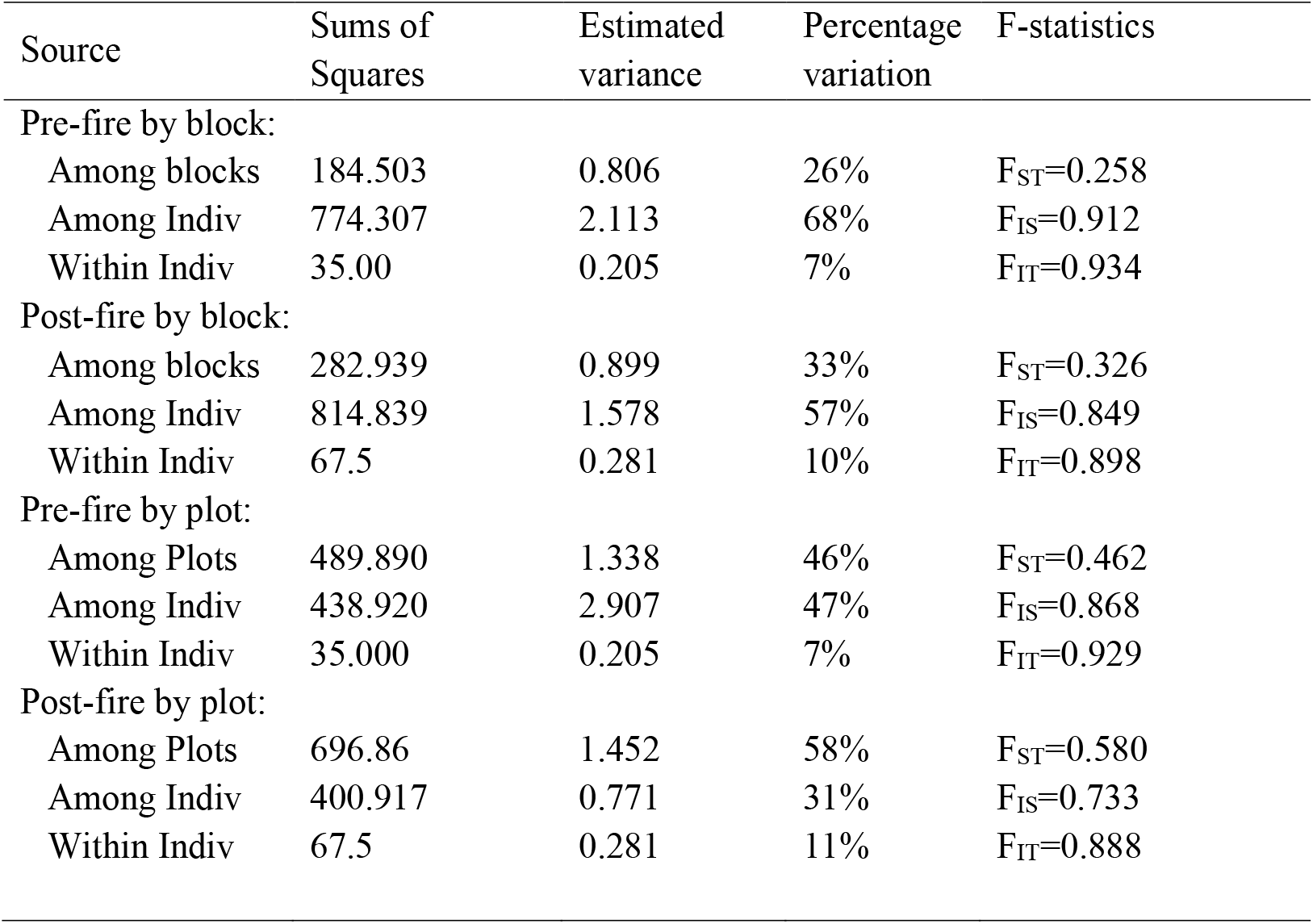
The effects of fire on the hierarchical partitioning of genetic variation in *Polygala lewtonii* using AMOVA analysis. Analyses were conducted before and after the 2016 fire, with individuals grouped by blocks, or plots.

When assessing changes in population structure independent of population designations using InStruct, genetic structure tended to increase after burning, with corresponding decreases in levels of admixture. The DIC and ln-likelihood values leveled out between K = 4 and K = 6 (Supporting Information–Figure S1), but we present K=5 in Figure 3 because it had the clearest assignment to clusters. Generally, a greater number of individuals were assigned to the most common genetic cluster in each plot or block, and individuals showed greater average assignments to the predominant cluster, indicating less admixture post-fire (Table 4). In eight out of 20 total plots (such as plots 15, 18, 25 and 28; Figure 3), the number of individuals assigned to the most common InStruct cluster increased post-fire, whereas it remained at 100% for eight plots, and decreased in only two plots. The proportion of admixture accounted for by the majority cluster also increased in all but two plots, indicating individuals were less admixed between InStruct groupings (Table 4). These results indicate that genetically similar individuals were more tightly spatially clustered after a fire and showed less admixture between plots.

**Table 4.** The effects of fire on genetic structure and admixture in *Polygala lewtonii*. Results show the proportion of individuals assigned to a majority cluster in InStruct pre- and post-fire, and the average admixture proportions of assignment to the predominant cluster in plots before and after the fire.

| Plot | Block | Pre-fire<br>proportion of<br>individuals<br>assigned to<br>the<br>predominant<br>cluster | Post-fire<br>proportion of<br>individuals<br>assigned to the<br>predominant<br>cluster | Pre-fire average<br>admixture<br>proportion for<br>predominant<br>cluster | Post-fire<br>average<br>admixture<br>proportion<br>for<br>predominant<br>cluster |
| --- | --- | --- | --- | --- | --- |
| 9 | 2 | 100% | 100% | 0.904 | 0.941 |
| 10 | 2 | 65.5% | 66.7% | 0.503 | 0.624 |
| 11 | 2 | 100% | 100% | 0.593 | 0.637 |
| 12 | 2 | 62.5% | 91.7% | 0.466 | 0.655 |
| 13 | 2 | 66.7% | 58.3% | 0.426 | 0.434 |
| 14 | 2 | 75.0% | 83.3% | 0.592 | 0.665 |
| 15 | 2 | 100% | 100% | 0.916 | 0.962 |
| 16 | 2 | 88.9% | 100% | 0.669 | 0.773 |
| 18 | 3 | 100% | 100% | 0.746 | 0.776 |
| 21 | 3 | 100% | 83.3% | 0.952 | 0.802 |
| 22 | 3 | 42.9% | 50% | 0.326 | 0.352 |
| 23 | 3 | 100% | 100% | 0.920 | 0.951 |
| 25 | 4 | 77.8% | 100% | 0.764 | 0.863 |
| 26 | 4 | 62.5% | 66.7% | 0.713 | 0.708 |
| 27 | 4 | 66.7% | 100% | 0.576 | 0.829 |
| 28 | 4 | 100% | 100% | 0.875 | 0.903 |
| 29 | 4 | 100% | 100% | 0.862 | 0.868 |
| 30 | 4 | 62.5% | 75% | 0.649 | 0.706 |
| 31 | 4 | 100% | 100% | 0.774 | 0.761 |
| 32 | 4 | 100% | 100% | 0.877 | 0.749 |
| Average<br>across<br>plots |  | 84% | 89% | 0.705 | 0.748 |

**Figure 3.**
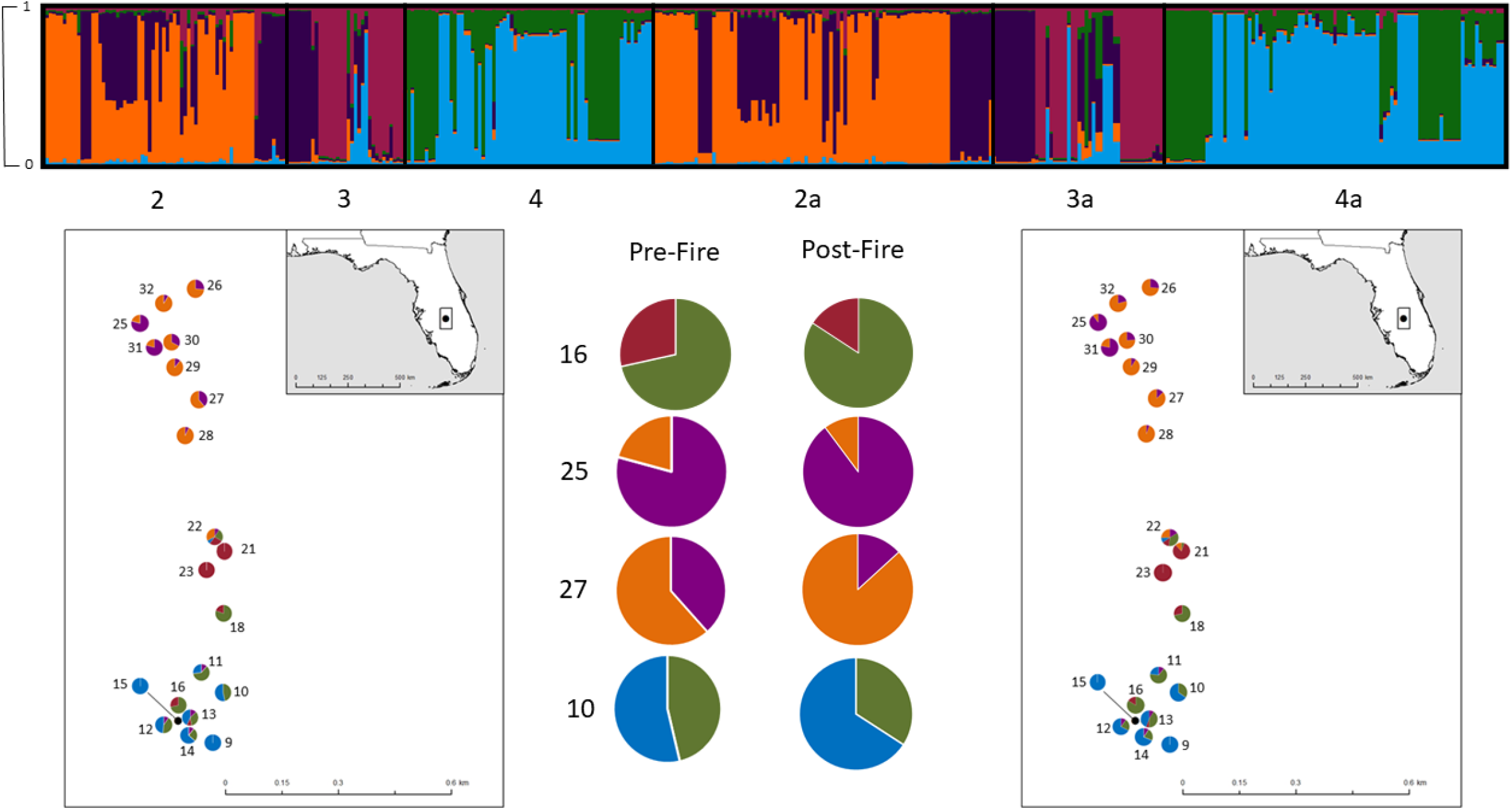
Patterns of genetic structure from pre- and post-fire populations show a decline in admixture as a result of fire. InStruct graph and maps showing population structure for pre- and post-fire, and corresponding pie charts. These cluster assignment proportions are also shown as pie charts superimposed on their respective sampling plots. The pie charts in the center (plots 16, 25, 27, and 10) are highlighted as they show examples of reduced post-fire admixture.

The results of analyses in which individuals were grouped by InStruct cluster were consistent with those in which individuals were grouped by block or plot. We observed minor changes in genetic diversity (Supporting Information–Table S3), an increase in among-group genetic variation and a decrease in within-group and within-individual variation in AMOVA analyses (Supporting Information–Table S4), and a decrease in admixture in InStruct (Supporting Information–Table S5) after a fire.

## Discussion

The overall goal of the study was to test the hypothesis that the role of seeds produced belowground in amphicarpic species is to recolonize a site after a disturbance (Cheplick and Quinn 1987, Cheplick and Quinn 1988). We focused on the amphicarpic species *P. lewtonii* and explored how genetic diversity and structure changed when the species recolonized a site after fire. Specifically, we tested whether we would find genetic patterns consistent with below-ground CL flowers primarily recolonizing a site after disturbance, or alternatively, whether genetic patterns would show that sites were predominantly re-colonized by the more prolific aboveground CH flowers, as proposed by Swift et al (2016). We quantified the outcrossing rate in aboveground CH flowers and genotyped pre- and post-fire samples, which allowed us to analyze shifts in genetic diversity, inbreeding, and genetic structure after fire.

To address the issue of differential germination after a disturbance, we first quantified outcrossing in seeds produced by aboveground CH flowers of *P. lewtonii* to determine whether it differed from that of CL flowers, which are obligately selfing. If the CH flowers relied fully on delayed selfing, then the seeds would be genetically indistinguishable from those produced by the CL flowers, in which case we would be unable to detect differential germination after a fire. Although 30 of 40 individuals (75%) had only maternal alleles present, indicative of selfing, the remaining 25% of individuals had at least one non-maternal allele, indicating reproduction via cross pollination. Although the selfing/inbreeding rate was still generally high in the aboveground CH flowers, simulations showed that the outcrossing rate differed significantly from that expected in a fully selfing mating system. The results of this experiment therefore indicate that it is possible to detect whether the aboveground CH flowers or belowground CL flowers predominantly germinated after a fire.

In the second experiment, we compared patterns of genetic diversity, inbreeding coefficients, and genetic structure in a natural population before and after a fire, results of which were most consistent with increased post-fire germination of seeds produced by belowground CL flowers. Observed and expected heterozygosity remained similar to pre-fire levels, but these values were already extremely low in pre-fire samples and were generally indicative of selfing. Inbreeding coefficients were very high pre-fire, but they increased further after fire. If seeds produced by aboveground CH flowers preferentially germinated after a fire, then given the greater average seed set of CH flowers (Koontz et al 2018) and the estimates of outcrossing obtained in the first experiment, we undoubtedly would have observed a post-fire increase in heterozygosity and a decrease in inbreeding coefficients. Given that this species is adapted to a disturbance-mediated ecosystem, this increase in inbreeding suggests that the species may rely on selfed seed as its primary means of recolonizing a site after fire.

Analyses of population structure also supported the conclusion that CL flowers preferentially germinated after a fire, and specifically showed a pattern consistent with the re-colonization of the site predominantly by seed produced by belowground CL flowers. Following fire, AMOVA analyses generally showed an increase in among-group variation (F_ST_) and a decrease in the variation found both among individuals within a group (F_IS_) and within individuals (F_IT_). These changes in the hierarchical partitioning of genetic variation are likely the result of post-fire germination by seeds that arose predominantly through selfing, which reduces effective population sizes and results in strong genetic drift, thereby resulting in strong genetic divergences among groups. This pattern was also found in InStruct results, which showed that genetically similar individuals were more tightly spatially clustered and that individuals showed lower admixture between genetic cluster after a fire. In general, greater among-population genetic structure arises from reduced gene flow, decreased dispersal distances and increased rates of inbreeding, all of which would most strongly occur in the seeds produced by the belowground CL flowers, which are obligately selfing and dispersed >1 m along the plant’s rhizome. Thus, changes in genetic structure observed after a fire support the hypothesis that the seeds produced by belowground CL flowers predominantly recolonized the site after fire. Although previous studies found that seeds buried belowground often showed greater germination after a fire (Cheplick and Quinn 1987, Cheplick and Quinn 1988), the current study is among the first to show that the plants that recolonized a site immediately after a disturbance most likely arose from seeds produced by the belowground CL flowers in an amphicarpic species. Additional studies are needed to see if this hypothesis is supported in other amphicarpic plants.

An increased contribution by seeds produced by belowground CL flowers seems especially plausible after a fire event, given that amphicarpy is hypothesized to be an adaptation ecological disturbance (Symonides, 1988). For amphicarpic species that inhabit fire-maintained ecosystems, belowground seeds may be over-represented in the seedbank from which the plant population germinates after fire because they occur deeper in the soil and may be less likely to be damaged by fire. This hypothesis is supported by previous research in another amphicarpic species, *Amphicarpaea bracteata*, that found that seeds sown at shallower depths (0-1 cm) had lower germination rates after fire than those sown deeper in the soil (Cheplick and Quinn 1987). The possibility that fire may damage aboveground seeds also helps explain why we did not find an increase in outcrossing after a fire, despite the fact that CH flowers produce much greater quantities of seed than either type of CL flowers (Koontz et. al. 2017). Another possible explanation for why the seeds produced belowground showed increased contributions to the population after a fire is because they experienced more favorable ecological conditions for germination than those produced aboveground (e.g., greater moisture below the soil surface). Indeed, previous studies in *Amphicarpaea* showed that after a fire, seedlings that arose from seeds sowed at shallower depths were more likely to desiccate than those sowed deeper in the soil (Cheplick and Quinn 1987). Future research in *P. lewtonii* is needed to understand the relative contribution of seed survival and environmental variables in promoting post-fire colonization of belowground seeds.

Overall, these results support a cyclical pattern to the reproduction, genetic diversity, and genetic structure in *P. lewtonii*, which appears to be an adaptation to a fire-maintained ecosystem. After a fire, the seeds from belowground CL flowers likely predominantly recolonize a site, resulting in low heterozygosity, low admixture, high inbreeding, and strong patterns of genetic structure immediately post-fire. In years that do not experience fire, most seedlings may arise from seeds produced by the more prolific aboveground CH flowers (Koontz et al 2018). Although the present study showed that only 25% the reproduction achieved by CH flowers was from outcrossing, several years of reproduction primarily arising from seeds produced by CH flowers would lead to a gradual increase in heterozygosity and a decrease in inbreeding coefficients and population structure, as was found in the pre-fire dataset. This cycle appears to be an adaptation to thrive in an environment that regularly experiences fire-related disturbance; the seeds produced belowground are protected from fire and may experience the most suitable conditions for germination immediately following a fire event, whereas the aboveground seeds produced by CH flowers may experience more suitable conditions for germination between fire events. This genetic variation generated by outcrossing in CH flowers could also help facilitate adaptation of *P. lewtonii* to environmental change (Oakley et al., 2007). Although many studies have debated the stability of mixed mating systems (Holsinger 1991; Goodwille et al 2005), the mixed mating system and associated amphicarpy in *Polygala lewtonii* and other amphicarpic species may be stable because it provides seeds that are well suited for germination in the contrasting environmental conditions found in disturbance-prone environments (Zhang et al 2020).

Interestingly, we observed a clear shift in alleles present in the population following the fire event, which provides some indication of seed longevity in the soil seed bank. Although some alleles were lost after fire, all of which were initially present in low frequencies within the population, making them vulnerable to loss through drift, a roughly equal number of new alleles emerged at low frequencies after fire. Of the alleles that emerged after fire, none were encountered in previous population genetic analysis of the species, even when samples from across *P. lewtonii’s* whole distribution were included in the analysis. Because this population was completely burned, seed dispersal is limited, and these alleles were not detected previously, it is likely that at least some of them were preserved in the soil seed bank and re-emerged after fire. Given that the first sampling occurred in 2014 and the alleles reappeared in 2017, this indicates that seeds may have remained viable in the seed bank at least four years, but possibly longer, before germination was stimulated by fire. Given that the success of the amphicarpic life history strategy depends on seeds maintaining viability in the soil, this provides further support for the hypothesis of post-disturbance recolonization by belowground seeds.

Finally, it is well known that fire maintains sandhill and scrub ecosystems and is an integral part of the natural cycle of disturbance (USFWS, 1999; Menges, 2007). Fire has multiple positive effects on *Polygala lewtonii*, including increased recruitment and increased survival of post-burn recruits (Weekly and Menges, 2012). This study highlights the highly specific adaptations that *P. lewtonii* has that allows it to thrive in response to fire and underscores the importance of using fire management to sustain populations of this endangered species (Weekley and Menges 2012). More generally, the results also support previous hypotheses about functional significance of amphicarpy as an adaptation to disturbance-prone environments. Given that amphicarpy has evolved independently multiple times in species occupying fire-maintained ecosystems (Zhang et al 2020), these results underscore the importance of fire as a selective force affecting the evolutionary history of plants as well as the importance of maintaining historical fire regimes to ensure the conservation of biodiversity.

## Acknowledgements

The authors thank Wendy Applequist, Monica Carlsen, and Peter Hoch for organizing the Missouri Botanical Garden Research Experience for Undergraduates program; the DNA analysis facility at Science Hill at Yale University for processing genetic samples; and Carl Weekley, Stephanie Koontz, Jamie Peeler, Devon Picklum, David Zaya, Jennifer Navarra, Sarah Hicks, Lauren Sullivan, John Lowell, Kate Prengaman, and David Horton for assistance with field work. This work was supported by the Florida Department of Agriculture and Consumer Services Division of Plant Industry (grant number 020159) and the Research Experiences for Undergraduates Program of National Science Foundation (grant number DBI-1157030).

