## Supplementary material for "Understanding how an amphicarpic species with a mixed mating system responds to fire: a population genetic approach": Table S1-5, Figure S1, Appendix S1

Table S1. Measures of individual heterozygosity (PHt, Hs-observed and expected, IR and HL) from genotyped seeds of *Polygala lewtonii* and simulated selfing. PHt, the proportion of heterozygous loci; Hs_observed and Hs_expected are standardized measures of heterozygosity, IR (internal relatedness) is a measure of heterozygosity which weights rare alleles more heavily, and HL is homozygosity by locus.

| Metric | Observed value in seeds | Mean values obtained under Simulation |
|---|---|---|
| PHt | 0.108 | 0.021 |
| Hs-Observed | 1.122 | 1.000 |
| Hs-Expected | 0.380 | 0.062 |
| IR | 0.680 | 0.935 |
| HL | 0.841 | 0.957 |

Table S2. Alleles unique to either the pre- or post-fire sample.

| ***Pre- or Post-fire*** | ***Locus*** | ***Allele*** | ***Frequency*** |
|---|---|---|---|
| Pre | PL56 | 194 | 0.006 |
| Pre | PL56 | 210 | 0.006 |
| Pre | PL80 | 134 | 0.006 |
| Pre | PL18 | 219 | 0.004 |
| Pre | PL18 | 246 | 0.008 |
| Pre | PL82 | 150 | 0.008 |
| Pre | PL54 | 131 | 0.006 |
| Post | PL40 | 165 | 0.004 |
| Post | PL40 | 175 | 0.004 |
| Post | PL80 | 152 | 0.031 |
| Post | PL80 | 156 | 0.021 |
| Post | PL18 | 234 | 0.007 |
| Post | PL18 | 244 | 0.027 |
| Post | PL82 | 173 | 0.004 |
| Post | PL82 | 175 | 0.013 |

Table S3. Genetic diversity in pre- and post-fire populations of *Polygala lewtonii* grouped by InStruct cluster. $N$, number of samples, $H_O$, observed heterozygosity, $H_E$, expected heterozygosity, $A$, average number of alleles per locus, $A_R$ allelic richness based on a minimum sample size of 21 individuals, $Ap$, number of private alleles, $F$, inbreeding coefficient, and $F^B$ the inbreeding coefficient taking null alleles into account,

| Instruct cluster | $N$ | $H_O$ | $H_E$ | $A$ | $A_R$ | $A_P$ | $F$ | $F^B$ |
|---|---|---|---|---|---|---|---|---|
| 1—Pre | 24 | 0.044 | 0.289 | 2.000 | 1.94 | 0 | 0.797 | 0.792 |
| 2—Pre | 37 | 0.034 | 0.286 | 2.455 | 2.22 | 1 | 0.868 | 0.843 |
| 3—Pre | 30 | 0.011 | 0.223 | 2.727 | 2.45 | 1 | 0.840 | 0.929 |
| 4—Pre | 46 | 0.037 | 0.214 | 2.455 | 2.14 | 3 | 0.808 | 0.808 |
| 5—Pre | 43 | 0.077 | 0.279 | 2.818 | 2.40 | 1 | 0.634 | 0.657 |
| Mean-Pre | | 0.041 | 0.258 | 2.491 | 2.23 | 1.2 | 0.789 | 0.806 |
| 1—Post | 22 | 0.037 | 0.252 | 2.000 | 1.87 | 0 | 0.705 | 0.852 |
| 2—Post | 52 | 0.021 | 0.315 | 2.818 | 2.56 | 2 | 0.946 | 0.925 |
| 3—Post | 36 | 0.046 | 0.248 | 2.909 | 2.49 | 0 | 0.751 | 0.825 |
| 4—Post | 67 | 0.030 | 0.204 | 2.273 | 2.06 | 0 | 0.883 | 0.841 |
| 5—Post | 63 | 0.107 | 0.291 | 3.364 | 2.46 | 2 | 0.581 | 0.598 |
| Mean-post | | 0.048 | 0.262 | 2.673 | 2.29 | 0.8 | 0.773 | 0.808 |

Table S4. Results of analyses of the partitioning of genetic variation in Polygala lewtonii using AMOVA. Analyses were conducted before and after the 2016 fire, with individuals grouped by InStruct clusters.

| Source | Sums of Squares | Estimated variance | Percentage variation | F-statistics |
|---|---|---|---|---|
| Pre-fire by Instruct clusters | | | | |
| Among Clusters | 257.165 | 0.846 | 28% | $F_{ST}$=0.281 |
| Among Indiv | 721.274 | 1.954 | 65% | $F_{IS}$=0.901 |
| Within Indiv | 38.500 | 0.214 | 7% | $F_{IT}$=0.929 |
| Post-fire by Instruct clusters | | | | |
| Among Clusters | 411.326 | 1.074 | 40% | $F_{ST}$=0.401 |
| Among Indiv | 686.451 | 1.320 | 49% | $F_{IS}$=0.824 |
| Within Indiv | 67.500 | 0.281 | 11% | $F_{IT}$=0.895 |

Table S5. The average admixture proportions of *Polygala lewtonii* individuals in InStruct clusters. Groupings were defined by the predominant InStruct assignment for each individual, before and after fire.

| InStruct cluster | Pre-fire average admixture proportion for majority cluster | Post-fire average admixture proportion for majority cluster |
|---|---|---|
| 1 | 0.838 | 0.947 |
| 2 | 0.866 | 0.851 |
| 3 | 0.837 | 0.875 |
| 4 | 0.888 | 0.881 |
| 5 | 0.847 | 0.915 |
| Average across clusters | 0.858 | 0.89 |

**Figure S1. Plots of** Deviance Information Criterion (DIC) and ln-likelihood (ln(K)) curves for InStruct analysis of Polygala lewtonii as presented in Fig. 3 at values of K=1-10. Analysis includes both pre- and post-fire samples. Values begin to plateau between K=4 and K=6.

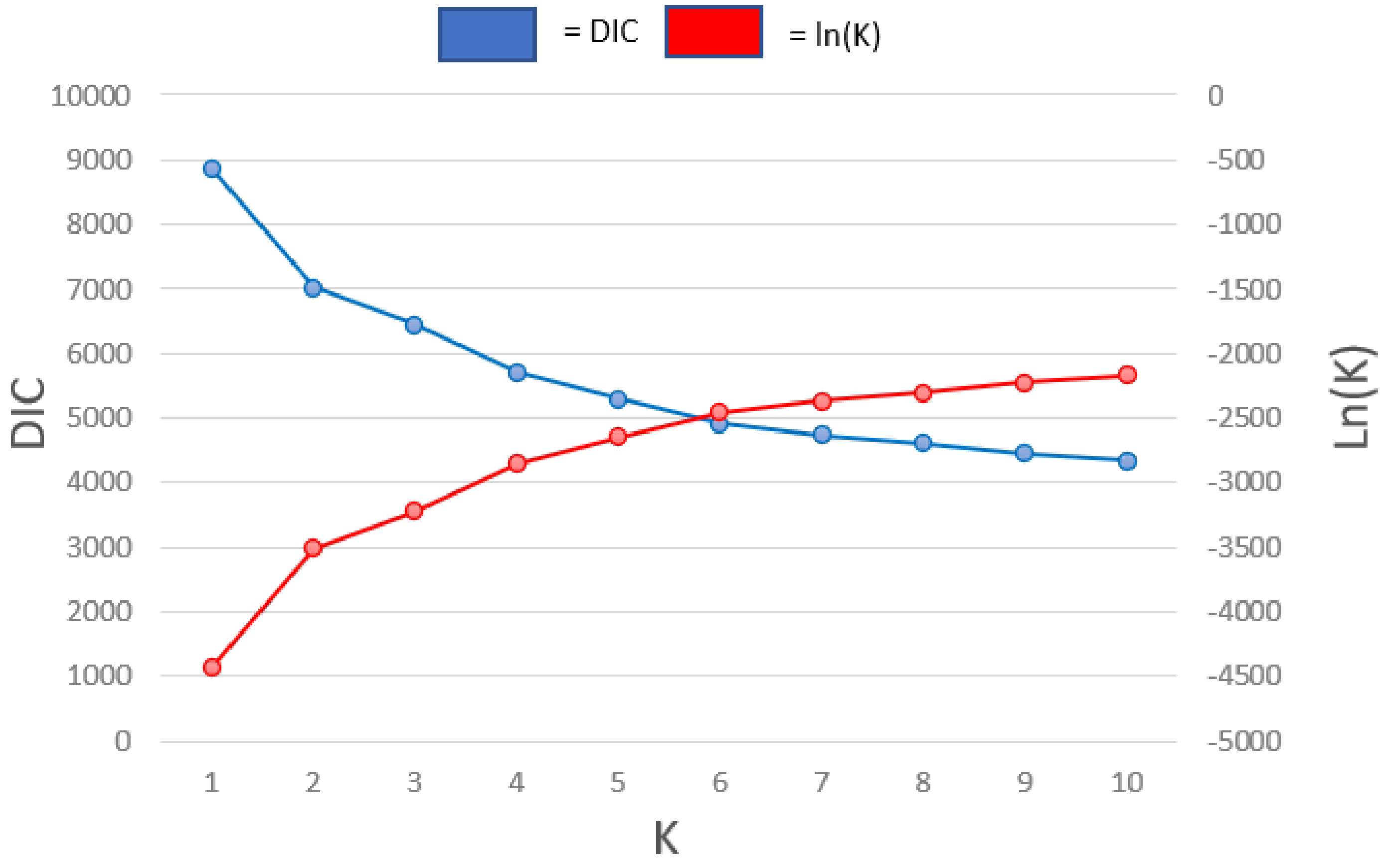

**Appendix S1**

Code for Inbreeding Simulation

```
setwd("U:/Scholar/Coauthor/Meyer/InbreedSim")
#install.packages("gtools")
library(gtools)
#reading the file
parents<-read.delim("ParentsGenotype.txt",sep="\t")
#generating the allele names
all<-unique(substr(names(parents)[-1],1,4))

#using sample to build 5 inbred seeds per parent with no LD
library(splitstackshape)

#bootstrapping the results 1000 times
#be sure to load GENEHET into the workspace before running this code

inbred.bootstrap<-c()

for(l in 1:1000){
inbred<-c()
for (i in 1:12){ #loci
for (j in 1:10){ #number of parents
#selecting loci by parent
loci<-unlist(c(parents[j,substr(names(parents),1,4)==all[i]]))
#pulling 10 gametes for 5 offspring
gametes<-sample(loci,10,replace=TRUE)
#adding the gametes into the set of seeds
inbred<-append(inbred,gametes,after=length(inbred))
}
}
dim(inbred)<-c(2,600) #reforming genotypes
genotypes<-paste(inbred[1,],inbred[2,],sep="_")
dim(genotypes)<-c(50,12)
colnames(genotypes)<-all
G<-data.frame(genotypes)
offspring<-
data.frame(paste0(rep(parents[,1],each=5),".",1:5),cSplit(indt=G,splitCols=names(G),sep="_"))
off.inbred<-GENHET(dat=offspring,estimfreq="T",locname=all)
num.inbred<-as.numeric(off.inbred[,2:6])
dim(num.inbred)<-c(50,5)
bootstrap.sample<-colMeans(num.inbred)
inbred.bootstrap<-
append(inbred.bootstrap,bootstrap.sample,after=length(inbred.bootstrap))
}
```

```
#reformatting the results of the bootstrapping into a column format
dim(inbred.bootstrap)<-c(5,1000)
bootstrapped.inbred.seeds<-t(inbred.bootstrap)
colnames(bootstrapped.inbred.seeds)<-colnames(off.inbred)[2:6]
write.csv(bootstrapped.inbred.seeds,"seeds.inbreeding.sim.csv")


#GENHET
#This program for R estimates 5 different individual heterozygosity estimates:
#proportion of heterozygous loci (PHt), standardized heterozygosity relative to the mean expected heterozygosity (Hs_exp, Coltman 1999),
#standardized heterozygosity relative to the mean observed heterozygosity (Hs_obs), IR (Amos 2001), HL (Aparicio 2006)


#parameters of the function:

#dat = matrix containing the genotypes; one row per individual; 1st column = individual identifier; following columns = the genotypes (1 allele (coded as a numeric value) per cell)
#missing data should be coded as "NA"
#see the example file provided with the code of the function, exGENHETgenotinput.txt

#estimfreq = binary variable taking the value "T" (true) or "F" (false): if estimfreq=T then the
#program will estimate the allele frequencies based on the data in the input file "dat"
#if estimfreq=F the program will use the allele frequencies provided by the user in another input file ("alfreq")

#locname = vector with the names of the different loci (in the same order as in "dat"); to provide only if estimfreq="T"

#alfuser = matrix containing the allele frequencies; to provide only if estimfreq="F";
#structure of alfuser: same number of columns as the number of loci,
#and same number of rows as the total number of alleles (over the different loci);
#each cell contains either the frequency of the allele or 0 (when, in the dataset,
#the locus considered does not have the allele considered); the first column contains
#the names of the alleles; the first row contains the names of the columns, i.e. "alleles"
#for the first cell, and then the names of the loci.

#alfuser can be produced by the function ALF, provided at the same web address as GENHET.

#if you import alfuser from a genalex output, MAKE SURE ALL THE DECIMALS ARE VISIBLE
```

```
	##########################################################################
#############
	##########################################################################
#############
	##########################################################################
#############
	##########################################################################
#############


	"GENHET"<-
	function(dat,estimfreq,locname,alfuser){

	nbloc=(ncol(dat)-1)/2
	nbind=nrow(dat)

	#estimation of allele frequencies (only if estimfreq=T)

	if(estimfreq=="T")

	{

	#creation of the list of alleles
	datv=vector(length=nbind*nbloc*2)
	for (i in 2:ncol(dat)) datv[(nrow(dat)*(i-2)+1):(nrow(dat)*(i-1))]=dat[,i]
	al=sort(na.omit(unique(datv)))

	#count of the number of times each allele appears + nb of missing data
	alcount=matrix(nrow=(length(al)+1),ncol=(nbloc+1))
	alcount[,1]=c(al,NA)
	for(j in 1:(nrow(alcount)-1))
		for(k in 1:(ncol(alcount)-1))
			alcount[j,(k+1)]=sum(dat[,(k*2):(k*2+1)]==alcount[j,1],na.rm=T)
	for(l in 2:ncol(alcount))
			alcount[nrow(alcount),l]=(2*nbind-sum(alcount[1:(nrow(alcount)-1),l]))


	#creation of the table of allele frequencies
	alfreq=matrix(nrow=length(al),ncol=(nbloc+1))
	colnames(alfreq)=c("Allele",locname)
	alfreq[,1]=al
	for(m in (1:nrow(alfreq)))
		for (n in 2:ncol(alfreq)) alfreq[m,n]=alcount[m,n]/(nbind*2-
alcount[nrow(alcount),n])

	}
```

```
else alfreq=alfuser


dat=as.data.frame(dat)
library(gtools)
res=matrix(nrow=nrow(dat),ncol=6)
colnames(res)=c("sampleid","PHt","Hs_obs","Hs_exp","IR","HL")
res[,1]=as.character(dat[,1])



#estimation of E per locus (for HL and Hs_exp)
E=vector(length=nbloc)
alfreq2=alfreq[,2:ncol(alfreq)]*alfreq[,2:ncol(alfreq)]
for(k in 1:ncol(alfreq2)) E[k]=1-sum(alfreq2[,k],na.rm=T)


#estimation of the mean heterozygosity per locus
mHtl=vector(length=nbloc)
ctNAl=0
ctHtl=0
for(l in 1:ncol(dat))
	{if (even(l)==T)
		{
		for (m in 1:nrow(dat))
			{ if (is.na(dat[m,l])==T) ctNAl=(ctNAl+1)
				else if (is.na(dat[m,(l+1)])==T) ctNAl=(ctNAl+1)
				 else if (dat[m,l]!=dat[m,(l+1)]) ctHtl=(ctHtl+1)
			}
		mHtl[l/2]=ctHtl/(nrow(dat)-ctNAl)
		ctNAl=0
		ctHtl=0
		}
	}

#the program in itself

ctHt=0
ctNA=0

ctHm=0
smHtl=0
mmHtl=0
sE=0
mE=0
```

```
sfl=0

sEh=0
sEj=0

for(i in 1:nrow(dat))
        { for (j in 2:(nbloc*2))
                { if (even(j)==T)
                        {
                        if (is.na(dat[i,j])==T) ctNA=(ctNA+1)
                        else if (is.na(dat[i,(j+1)])==T) ctNA=(ctNA+1)
                           else {
                                      if (dat[i,j]!=dat[i,(j+1)])
                                              {
                                               ctHt=(ctHt+1)
                                               sEj=sEj+E[j/2]
                                              }
                                     else sEh=sEh+E[j/2]
                                     smHtl=smHtl+mHtl[j/2]
                                     sE=sE+E[j/2]

        sfl=sfl+alfreq[alfreq[,1]==as.numeric(dat[i,j]),(j/2+1)]+alfreq[alfreq[,1]==as.numeric(dat
[i,j+1]),(j/2+1)]
                                     }
                        }
                }
         res[i,2]=ctHt/(nbloc-ctNA)
         mmHtl=smHtl/(nbloc-ctNA)
         res[i,3]=(ctHt/(nbloc-ctNA))/mmHtl
         mE=sE/(nbloc-ctNA)
         res[i,4]=(ctHt/(nbloc-ctNA))/mE
         ctHm=nbloc-ctHt-ctNA
         res[i,5]=(2*ctHm-sfl)/(2*(nbloc-ctNA)-sfl)
         res[i,6]=sEh/(sEh+sEj)
         ctHt=0
         ctNA=0
         ctHm=0
         smHtl=0
         mmHtl=0
         sE=0
         mE=0
         sfl=0
         sEh=0
         sEj=0
         }
```

```
	return(res)
	}

	################################################################################
##############
	################################################################################
##############
	################################################################################
##############
	################################################################################
##############
```
